# Dynamic interactions of the coronavirus PEDV nucleoprotein with its viral partner nsp3 and RNA

**DOI:** 10.64898/2026.11.24.769051

**Authors:** Cécile Fétré, Julie Groutsch, Magali Aumont-Nicaise, Philippe Fossé, Olivier Mauffret, Quentin Nevers, Bernard Delmas

## Abstract

While coronavirus genome replication and transcription occur inside double-membrane vesicles (DMVs) in infected cells, nucleocapsid assembly of genomic RNA takes place at their cytoplasmic (external) face. Neosynthesized viral mRNAs (including genomic RNA) are exported from the DMV through nsp3/nsp4 pores and genomic RNAs are believed to be selected for encapsidation through an essential interaction of the nucleoprotein N with the amino-terminal domain of nsp3. Here, we describe the interaction play between the N of porcine epidemic diarrhea virus (PEDV, an alphacoronavirus) with the N-terminal domain of nsp3 and with RNAs. While N interacts with the N-terminal domain of nsp3 with a dissociation constant (K_D_) of about 150 nM, it binds to diverse small RNAs, displaying an affinity in the pM range for a short RNA (RNA[8mer]). This RNA and a 60-nt long RNA hairpin were both able to fully displace nsp3 complexed with N. In contrast, nsp3 was not able to dissociate RNA from N-RNA complexes. Using truncated forms of N and of the N-ter domain of nsp3, we showed that the RNA binding site does not fully overlap the nsp3 binding site on N. These results suggest that the interaction play between N, nsp3 and RNA is unidirectional, allowing to propose a mechanistical model for genome encapsidation, model in which N is first recruited by nsp3 at the cytoplasmic face of the DMV to bind to nucleotide stretches of (genomic) RNA exiting from the nsp3/nsp4 channel and concomitantly to dissociate from nsp3.

## Introduction

With the recurrent emergence and re-emergence of coronaviruses in humans and economically important animal species, this group of viruses represent a main burden in both public health and livestock production sector. Although vaccines are currently used to fight against human coronaviruses such as SARS-CoV-2 that leads to the COVID-19 pandemic and animal coronaviruses, mainly in pig and poultry industries (Wang et al., 2019), it is necessary to better understand the functioning of the coronavirus replication cycle to identify new viral vulnerabilities for the design of antiviral strategies.

While the three highly-pathogenic human coronaviruses that emerged in human in the last 25-years (the first two being SARS-CoV in 2002 and Middle East respiratory syndrome coronavirus (MERS-CoV) in 2012) are representatives of the *β-coronavirus* genus in the family *Coronaviridae*, several pig coronaviruses with strong economic impact such as porcine epidemic diarrhea virus (PEDV) and transmissible gastroenteritis virus (TGEV) are phylogenetically distant from these three human viruses and belong to the *α-coronavirus* genus. Human coronaviruses 229E and NL63 and animal coronaviruses like feline infectious peritonitis virus, canine coronavirus and numerous bat coronaviruses also belong to this last coronavirus genus (Millet et al., 2021).

Coronaviruses are enveloped positive-sense single strand RNA viruses that express their own replication machinery. Replication and transcription of their genome (which is approximatively 30 kb in size) is carried out by RNA-dependent RNA polymerase complexes (RTCs, whose functioning has been partially elucidated by cryo-electron microscopy [Wang et al., 2020; Yan et al., 2021a; Yan et al., 2021b; Anderson et al., 2024), that derived from the proteolytic process of two large overlapping polyproteins, pp1a and pp1ab encoded by its 5’-terminal two-third part (V’kovski et al., 2021). RTCs are associated with membrane organelles called double-membrane vesicles (DMVs) generated from membranes deriving from the host secretory pathway (Wolff et al., 2020 and references therein). DMVs have been shown to constitute active sites for viral RNA synthesis (Neuman et al., 2014; Snijder et al., 2020).

In infected cells, multiple copies of the nucleoprotein (N) encapsidate the viral genome, providing protection from the host cell environment (Masters, 2026). N colocalizes to the RTCs of murine hepatitis virus (MHV, another β-coronavirus) and SARS-CoV-2, a feature that depends on an essential interaction with the N-terminal component of nonstructural protein nsp3 (Hurst et al., 2013; Cong et al., 2020). Viral genome-N complexes are found in ERGIC compartment where virus particles budding occurs (Klumperman et al., 1994; Krijnse-Locker et al., 1994; Ghosh et al., 2020).

Taking the SARS-CoV-2 N as a prototypal coronavirus nucleoprotein, it was shown that it is highly dynamic, made of three intrinsically disordered regions (IDRs, numbered N1, N3 and N5 constituting roughly 55% of the N length), intercalated with two folded domains N2 and N4 of about 130 and 100 residues, respectively (Bessa et al., 2022). N self-assembles into dimers in solution, is highly basic, and RNA binding is through to be cooperative and to associate all the five domains of N, playing a role in both RNA chaperoning and packaging (McBride et al., 2014; Zuniga et al., 2007; Zargarian et al., 2025). Electron tomography of SARS-Cov-2 virions identified numerous N-RNA complexes inside the SARS-CoV-2 virions, apparently associated with different domains of the genomic RNA. While the SARS-CoV-2 N3 IDR and of all other coronaviruses contains a serine-arginine (SR)-rich domain, its PEDV homolog displays downstream this domain a partially overlapping Asn-rich stretch (**Fig. 1**). Pairwise alignment between the N of SARS-CoV-2 and PEDV only revealed 43 and 30% amino acid identity between their N2 and N4 domains, and no match between their N1, N3 and N5 domains. However, AlphaFold3 structure predictions carried out on the N of PEDV as well as the X-ray structure of its N4 domain (PDB 8WQK) suggests that it may exhibit similar structure than its SARS-CoV-2 homolog.

**Figure 1:**
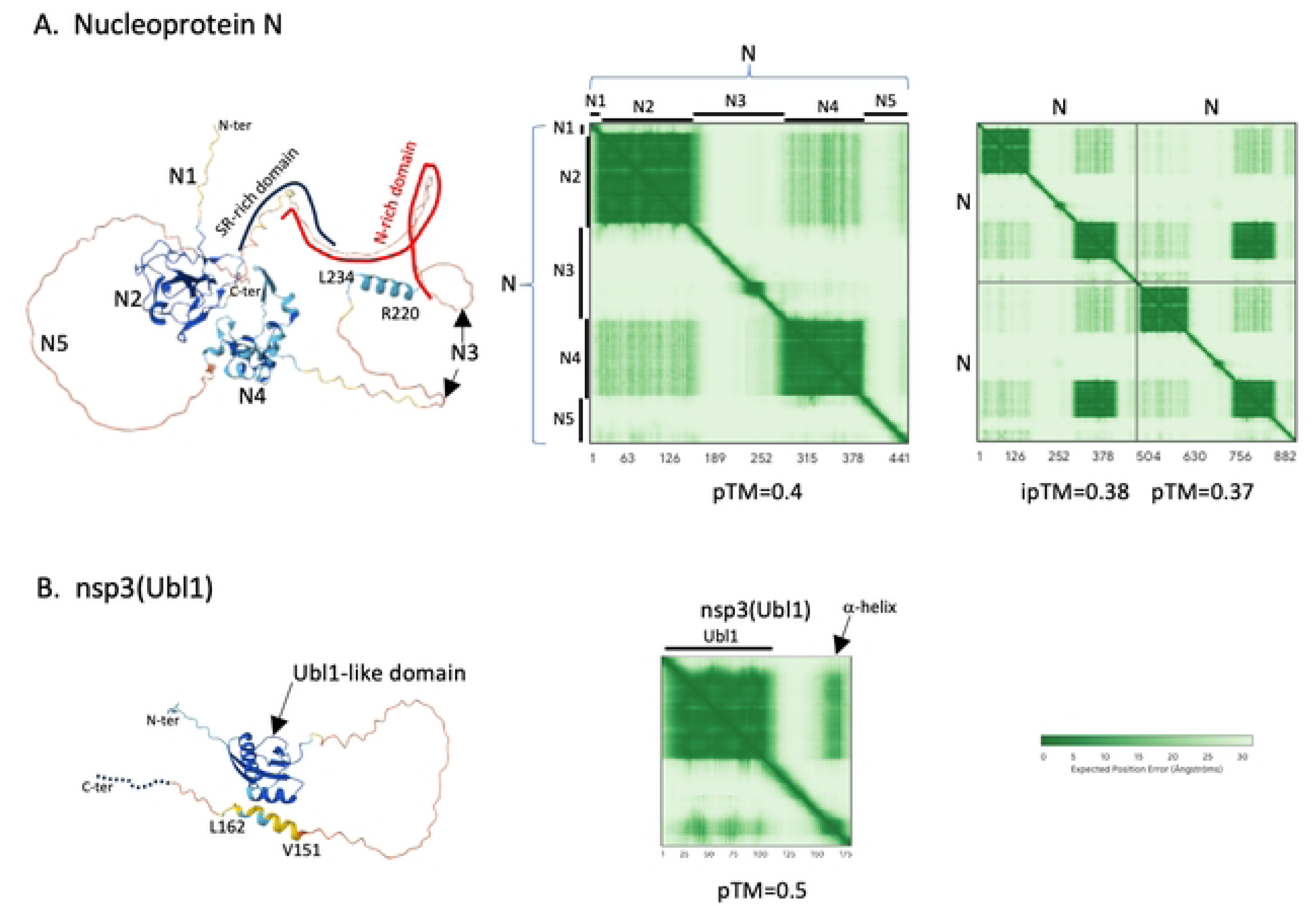
AlphaFold3 predicted structures of the PEDV nucleoprotein N (A) and the N-ter domains of nsp3 (B). On the left, AF3 structure predictions. On the right, the expected position error (in Angströms) of the best predicted structure of N as a monomer and a dimer, and the N-ter domains of nsp3. Note the N4 dolmain predicted as implicated in dimerization.

The non-structural protein 3 (nsp3), a critical N cofactor, is the largest protein encoded by coronaviruses, comprising roughly 2,000 amino acids, constituting up to 16-18 distinct domains and playing an essential role in coronavirus replication and transcription (Neuman, 2016; Lei et al., 2018, Huang et al., 2024). The domain organization is variable in different CoV genera. Most nsp3 domains (NH_2_-Ubl1-Mac1-Mac2-Mac3-DPUP-Ubl2-PL2pro-NAB and Y1-CoV-Y-COOH for β-coronaviruses of the clade B) are localized in the cytoplasmic side of the pore, constituting the cytoplasmic ring (Lei et al., 2018). However, in α-coronaviruses, an additional protease, PL1pro is encoded downstream the Ubl1 reading frame. Nsp3 and nsp4 are the minimal viral components required to induce DMV formation and to form a double-membrane-spanning pore, essential for the transport of newly synthesized viral RNAs outside the DMVs (nsp3-nsp4 pore complex comprises 12 copies of each of nsp3 and nsp4, (Wolff et al., 2020; Huang et al., 2024).

The first nsp3 amino-terminal domain that comprises an ubiquitin-like domain (Ubl1) was shown being involved in binding to the serine-rich (SR) region of N of MHV (Hurst at al., 2010), while a yeast two-hybrid screening identified a hydrophobic linear motif slightly upstream of the SR region (Cong et al., 2020). As determined by NMR and X-ray for SARS-CoV-2, the interaction between nsp3 and N proteins was shown to implicate two linear motifs in the N3 linker, a hydrophobic helix (residues 219 to 230), a disordered polar strand (residues 243 to 255) as well as the N2 structured domain (Bessa et al., 2022; Ni et al., 2023).

Here, we used electrophoretic mobility shift assays (EMSA) to describe the interaction of N of an α-coronavirus, PEDV, with the N-terminal domain of nsp3 and various RNA species. We show that the N-nsp3 interaction through its Ubl1 domain is mediated by the N-ter domains of N. Furthermore, we showed that small RNAs associate with N affinities in the picomolar-range and were able to dissociate the N-nsp3 complex, allowing RNA recruitment by N. In contrast, nsp3 was unable to displace the RNA bound to the N proteins, suggesting that nsp3 is involved in trafficking and chaperoning N before genome encapsidation.

## Results

### EMSA identifies PEDV N, but not nsp3(Ubl1), as a RNA binder

Interaction between recombinant purified N or nsp3(Ubl1) (**Fig. S1**) with short RNAs (7-to 11-nucleotides long) was investigated using EMSA of proteins migration on agarose native gels (**Fig. S2**). Coincubation of N with RNAs led to the identification of a single band specific to each N-RNA complex. While the N complexed with the RNA-7mer and RNA-11mer display a diffuse band in the native gel, N interacting with the RNA-8mer migrates as a discrete band, suggesting that N-RNA-8mer complexes are more homogeneous than with the two other RNAs. We noticed that N without RNA does not migrate substantially in the native gel to reside at the well - agarose gel interface. In the same experimental conditions, nsp3(Ubl1) was not able to bind the RNAs. Thus, the RNA-8mer was used for further N-RNA interaction analyses. To further characterize the interaction between N and RNA-8mer, we showed that N-RNA complexes formation is dependent of the concentration of N (**Fig. 2A**). As a negative control, nsp3(Ubl1)-RNA complex formation was not observed regardless of its concentration. Finally, we showed that N is able to interact with both RNA-8mer and its homologous DNA-8mer oligonucleotide (**Fig. S3**).

**Figure 2:**
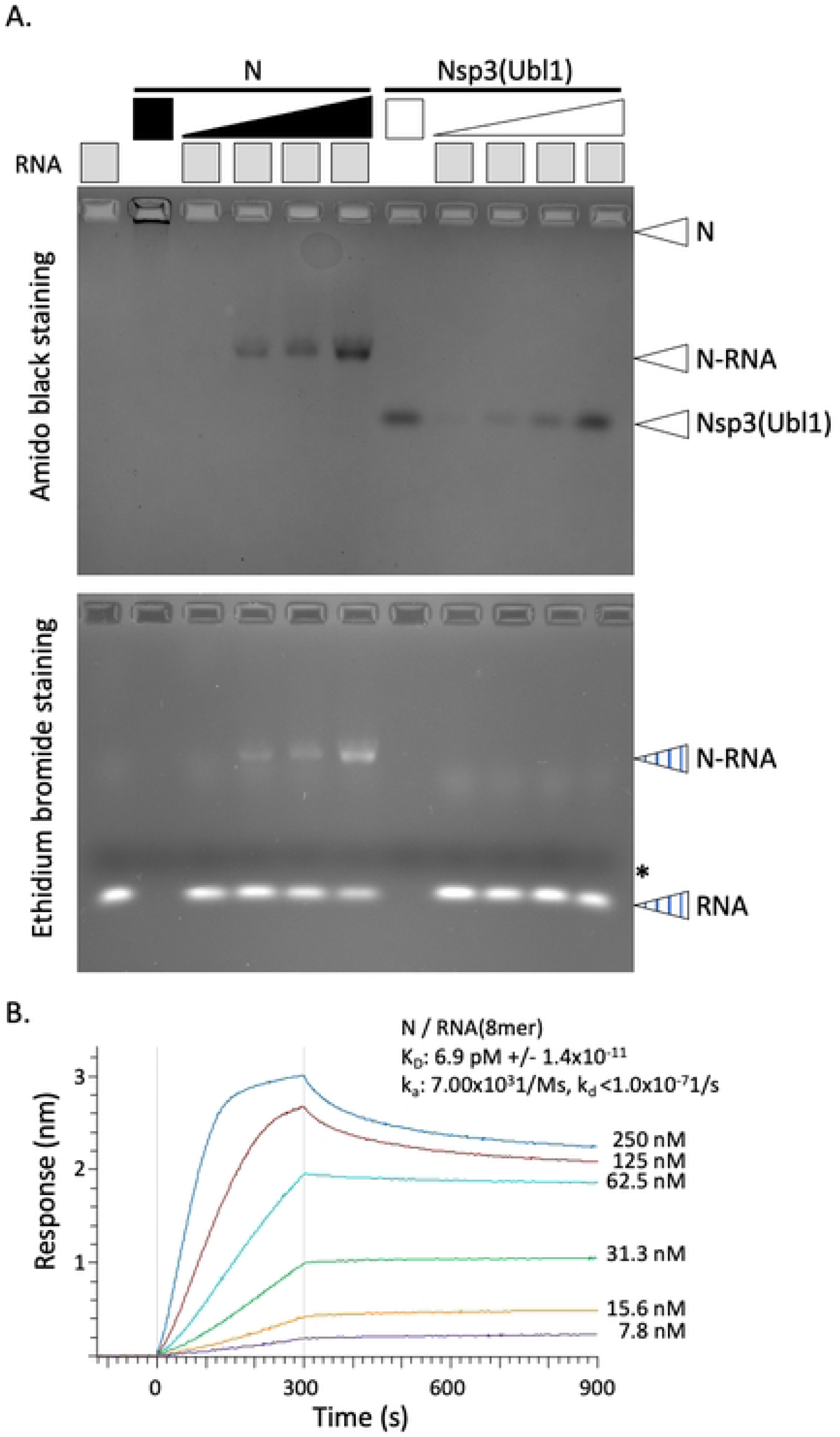
Respective abilities of N and nsp3(Ubl1) to bind RNA(8mer) oligonucleotide. (A) Native agarose gel electrophoresis of individual proteins N and nsp3(Ubl1) and their RNA-complexes. The gel was stained with Amido black to reveal proteins (upper panel) and with ethidium bromide for nucleic acids detection (lower panel). Bands corresponding to the proteins, free RNA and RNA-protein complexes are indicated. N and nsp3(Ubl1) concentrations were 0.5, 1, 2 and 5 mM (and 5 mM in the absence of RNA). RNA(8mer) concentration was 25 mM. * indicates the bromophenol blue tracking dye migration. (B) BLI binding kinetics measurements are shown for N to a biotinylated RNA. Equilibrium dissociation constants (KD) were determined on the basis of fits, applying 1:1 interaction model; ka, association rate constant; kd, dissociation rate constant.

We next explored the affinity of N for a biotinylated form of the RNA-8mer used as a bait by biolayer interferometry (BLI). The dissociation constant (Kd) value was far below 1 nM, which is too strong to measure reliably affinity with this technique (**Fig. 2B**). Association and dissociation curves were found compatible to a 1:1 interaction model.

### PEDV N forms stable complexes with nsp3(Ubl1)

Since the N and nsp3 of the two β-coronaviruses MHV and SARS-CoV2 were found to interact during infection, we aimed at determining if their α-coronavirus PEDV homologs are able to bind. An His-tagged version of the PEDV N was expressed in *E. coli* and purified to serve as a bait in BLI experiments (**Fig. 3A**). We then explored its affinity for a purified PEDV nsp3(Ubl1) form at different concentrations to determine its kinetic rate constants. Its affinity for nsp3(Ubl1) was about 150 nM, suggesting that PEDV N - nsp3(Ubl1) complexes may be stable at relatively high concentrations (**Fig. 3B**). Thus, we aimed at producing a PEDV N-nsp3(Ubl1) complex to delineate the interaction mode of N with its two ligands, RNA and nsp3. For this purpose, we co-expressed a GST-tagged form of nsp3(Ubl1) with the PEDV N, purified the complex by GST-tag affinity chromatography and followed by TEV cleavage and gel filtration chromatography. **Fig. 3C** shows that the PEDV N - nsp3(Ubl1) form a stable complex that was purified at nearly homogeneity.

**Figure 3.**
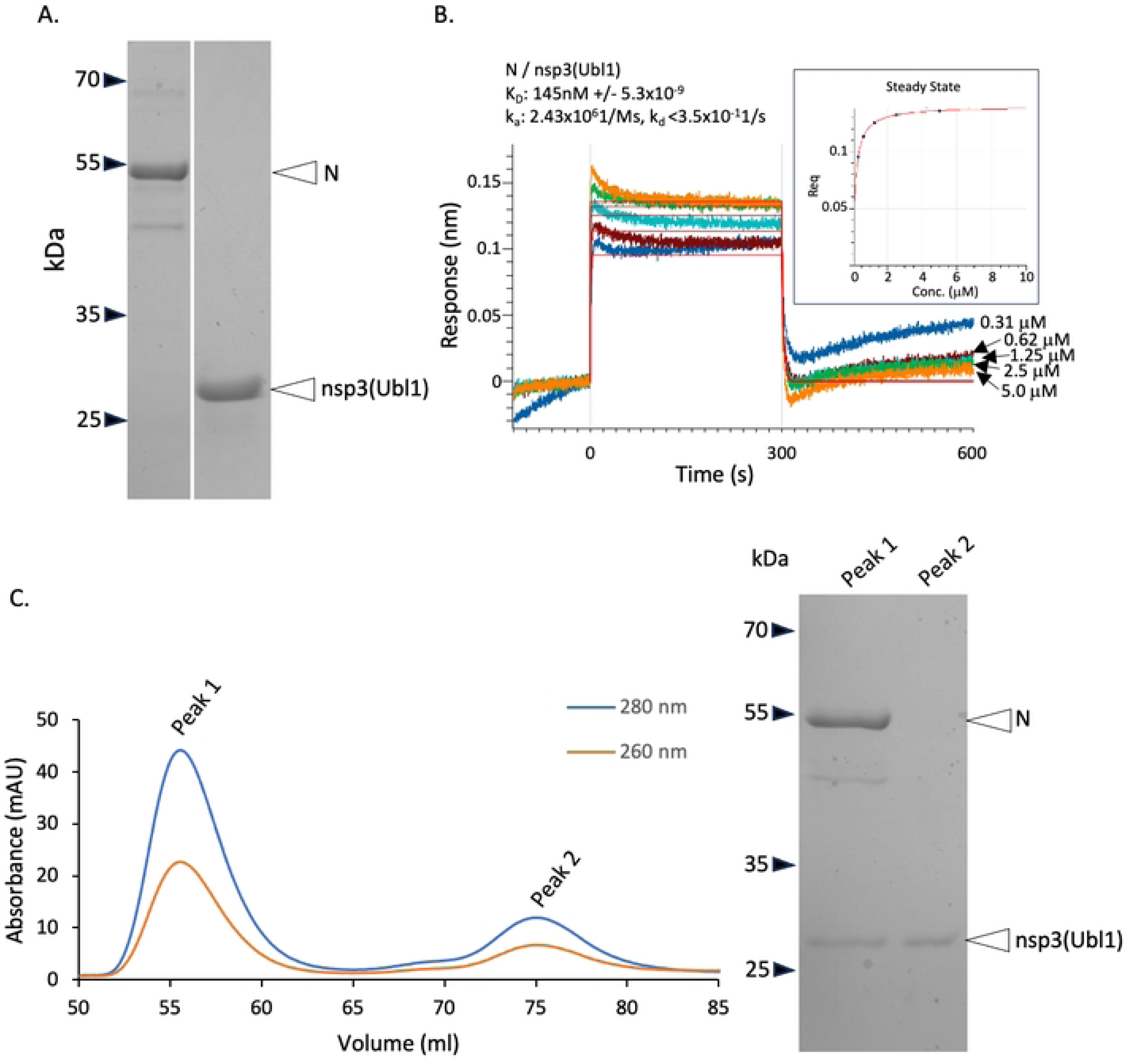
N-nsp3(Ubl1) affinity and complexe purification. (A) SDS-PAGE of purified PEDV N and nsp3(Ubl1) proteins. (B) BLI experiments to determine N – nsp3(Ubl1) affinity. (C) The N and the GST-nsp3(Ubl1) fusion protein were co-expressed in E. coli and submitted to affinity chromatography using glutathion agarose beads. Agarose-bound N / GST-nsp3(Ubl1) complexes were digested by the TEV protease to release N-nsp3(Ubl1) complexes. N - nsp3(Ubl1) complexes were pooled and fractionated by gel filtration chromatography (left). Two peaks were identified. Right, coomassie blue staining of a SDS-PAGE revealing peaks 1 and 2 protein contents. N and nsp3(Ubl1) form a stable complex with an KD of about 150 nM.

### RNA displaces nsp3 from N-nsp3 complexes to bind to N

To investigate the interplay between these three interacting species, RNA, N and nsp3 proteins, we carried in parallel two EMSA assays, a first one in which N was first complexed with RNA(8mer) before incubation with different concentrations of nsp3(Ubl1) (**Fig. 4A)**, and a second in which purified N:nsp3(Ubl1) complexes were incubated with increasing amount of RNA(8mer) (**Fig. 4B)**. Next, the composition of the resulting complexes was analyzed in the two assays, proteins being labeled by amido black staining and RNA by ethidium bromide. **Fig. 4A** shows that in all tested conditions (even with a nsp3:N ratio of 2), nsp3 was not able to bind to N complexed with the RNA. Furthermore, the N:RNA ratio remains constant whatever the nsp3 concentrations. No N-nsp3-RNA complex was evidenced. We concluded that nsp3 binding site on N may be masked by bound RNA or that N bound to RNA does not adopt a conformation suitable for nsp3 binding. In contrast, in the second assay, **Fig. 4B** shows that RNA(8mer) is able to displace nsp3(Ubl1) complexed with N in a concentration-dependent manner, and concomitantly to bind to N to form a stable N-RNA complex. No trace of RNA was evidenced on the nsp3 species excluded from the N-nsp3 complexes. To determine if the ability of the RNA to displace nsp3 from N-nsp3 complexes and to bind N is dependent of the nature of the nucleic acid, RNA or DNA, purified N:nsp3(Ubl1) complexes were incubated with increasing amount of DNA(8mer) (**Fig. S4)**. As found with RNA(8mer), its single-strand DNA homolog was able to displace nsp3 from the N-nsp3 complex and to bind to N. To summarize, these results show that N-nsp3 complexes can recognize (small) oligonucleotides, and that this binding results in the release of nsp3 previously complexed with N to form N-RNA complexes. These results also appear in full accordance with the difference of affinities of N to nsp3 and RNA, which are of the nM-μM range for nsp3 and the pM range for RNA.

**Figure 4:**
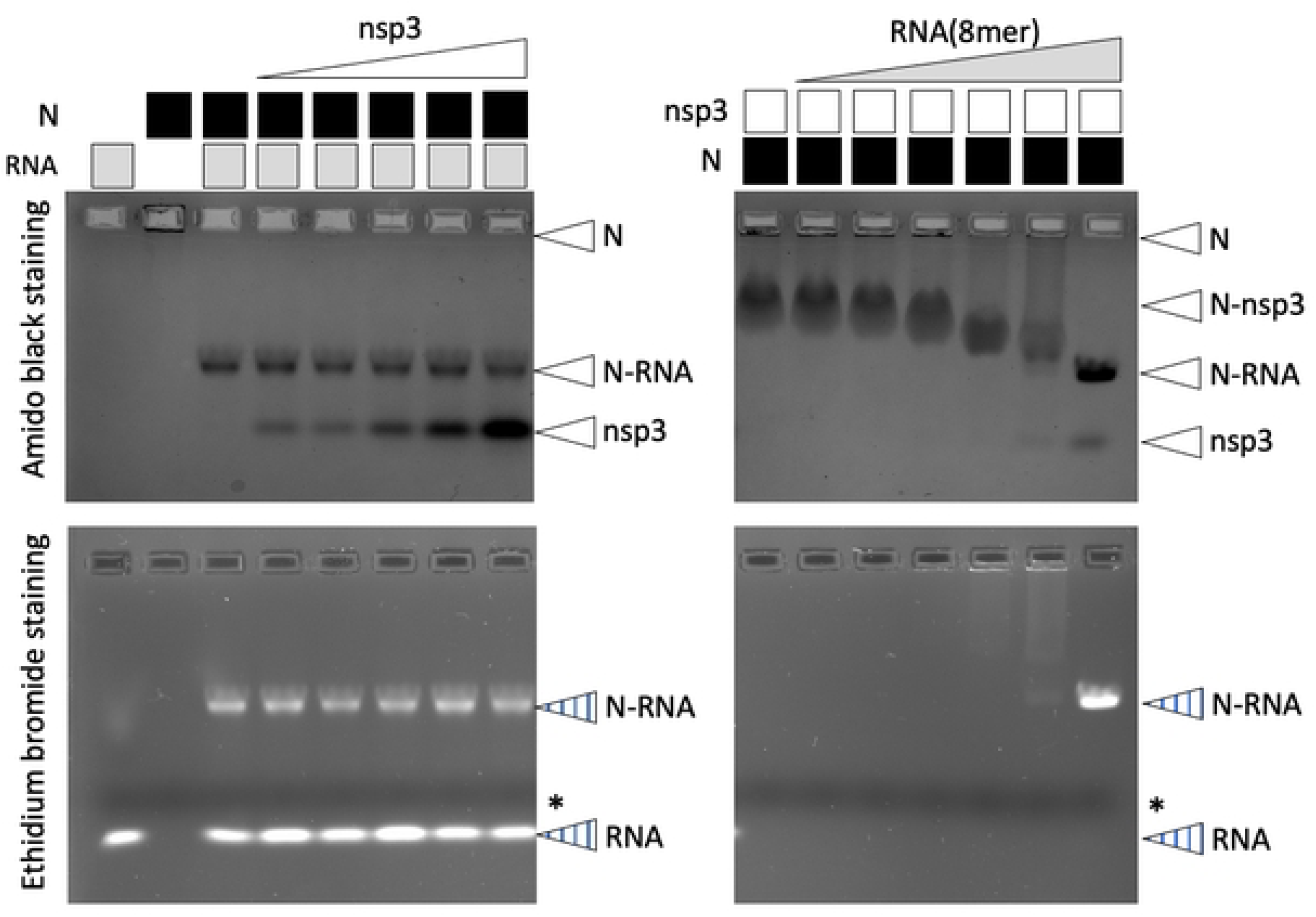
RNA displaces nsp3 from the N-nsp3 complex to associate to N. Left, native agarose gel electrophoresis of N-RNA(8mer) complexes (at concentration of 5 mM) incubated with increasing amounts of nsp3 (0, 1, 2, 5, 10 and 20 mM). Right, native agarose gel electrophoresis of the N-nsp3 complexe (at a concentration of 5 mM) incubated with increasing amounts of RNA(8mer) (0, 0.1, 0.5, 1, 2, 5 and 10 mM). The gels were processed as in Fig. 3. Bands corresponding to the individual proteins N and nsp3, N-nsp3 and N-RNA(8mer) complexes, and unbound RNA(8mer) are indicated. * indicates the bromophenol blue dye migration. While RNA(8mer) displaces nsp3 from the N-nsp3 complex, nsp3 is not able to displace RNA(8mer) from the N-RNA complexes.

### Hairpin RNA displaces nsp3 from PEDV N-nsp3 complexes to bind to N

The N protein of SARS-CoV-2 has been reported to act as an RNA chaperone protein able to bind and destabilize nucleic acid secondary structure and anneal nucleic acid stands (Zargarian et al., 2025). We next investigated if a hairpin RNA (hpRNA) can also be recognized by the PEDV N-nsp3 complex, resulting to its destabilization and to the binding of the RNA to the N protein. **Fig. 5, right panel** shows that, as found with RNA(8mer), hpRNA destabilizes the N-nsp3 complex, resulting to its binding to N and the release of nsp3. In contrast, nsp3 was not able to displace the N-hpRNA complex (**Fig. 5, left panel).**

**Figure 5:**
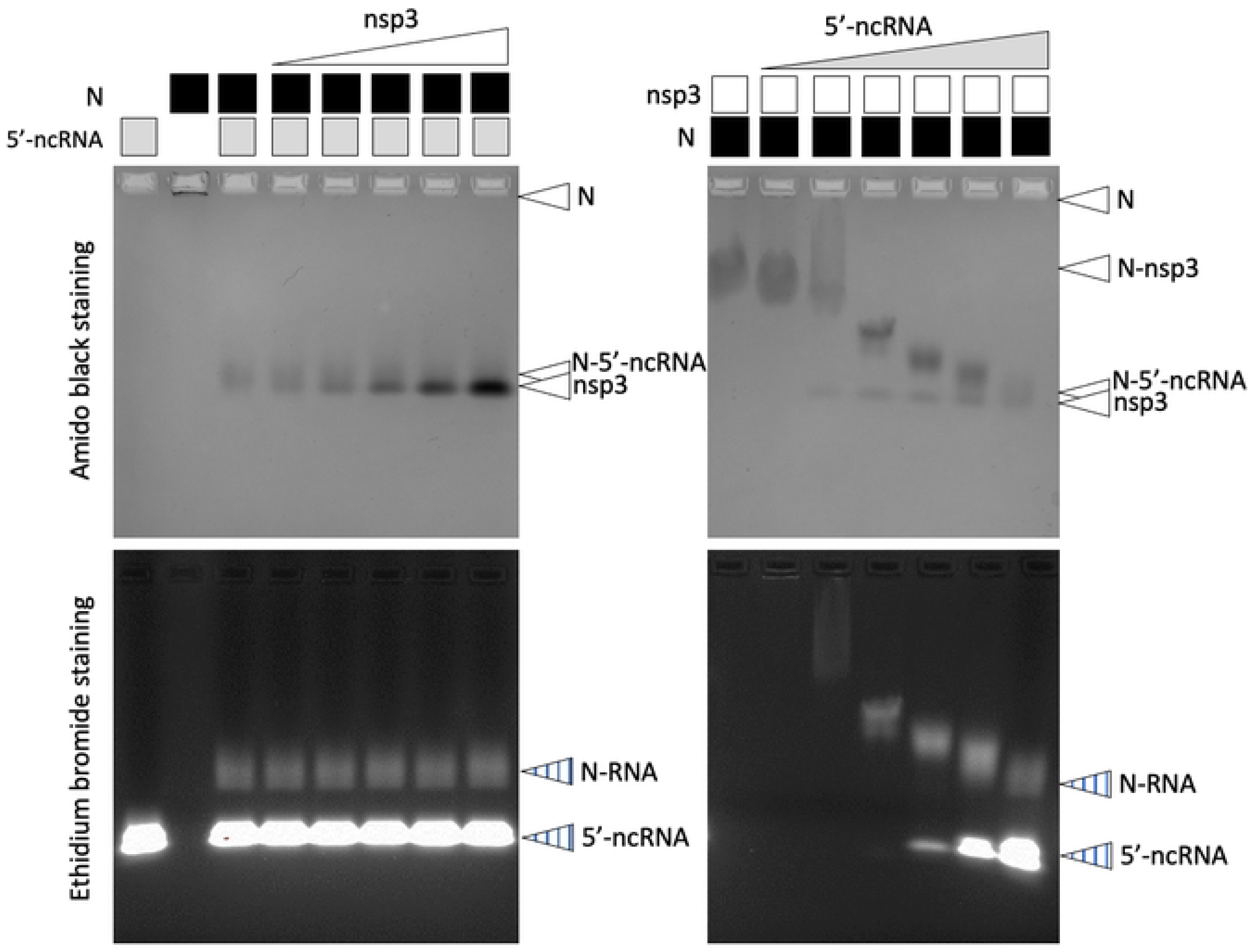
Hairpin RNA displaces nsp3 from the N-nsp3 complex to bind N. Left, native agarose gel electrophoresis of N-RNA(5’-ncRNA) complexes (at concentration of 5 mM for N and 15 mM for RNA(8mer)) incubated with increasing amounts of nsp3 (0, 1, 2, 5, 10 and 20 mM). Right, native agarose gel electrophoresis of the N-nsp3 complexe (at a concentration of 5 mM) incubated with increasing amounts of RNA (5’-ncRNA) (0, 0.1, 0.5, 1, 2, 5 and 10 mM). The gels were processed as in Fig. 3. Bands corresponding to the individual proteins N and nsp3, N-nsp3 and N-RNA(5’-ncRNA) complexes, and unbound RNA (5’-ncRNA) are indicated. While large RNA displaces nsp3 from the N-nsp3 complexe, nsp3 is not able to displace RNA from the N-RNA complexes.

### nsp3(Ubl1) and RNA binding sites on N are distinct

To further characterize the interplay between RNA and nsp3 and their exclusive binding to N, we used AlphaFold3 to predict their respective binding sites. **Fig. 6A** shows that the Ubl1 domain of nsp3 is expected to bind two distinct domains of N, its N2 folded domain and an α-helix present in the largely unfolded domain N3. RNA(8mer) is predicted to bind to the two N structured domains N2 and N4. Two plasmid constructs were generated, a first encoding the ubl1 domain of nsp3 (named Ubl1) and a second encoding a truncated form of N (called sN for “short-N”) in which a minimal linker was conserved between N2 and the N3 α-helix and the residues downstream the N3-helix deleted. sN and Ubl1 were predicted to interact by AF3 with an ipTM score of 0.85 (**Fig. 6B**). These two constructs were co-expressed in *E. coli* and purified as a complex through a GST-tag fused to Ubl1. The Ubl1-sN complex was released after TEV cleavage and submitted to gel filtration for purification (**Fig.6C**). SDS-PAGE analysis shows that the sN-Ubl1 complex is purified to homogeneity. The purified Ubl1-sN complex was incubated to increasing amounts of RNA(8mer) to quantify its ability to release Ubl1 from the complex (**Fig.6D**). In contrast to what was observed with the full-length N complexed with nsp3(Ubl1), RNA(8mer) was not able to displace Ubl1 from its complex with sN, suggesting that nsp3 and RNA(8mer) binding sites partially overlap on N.

**Figure 6:**
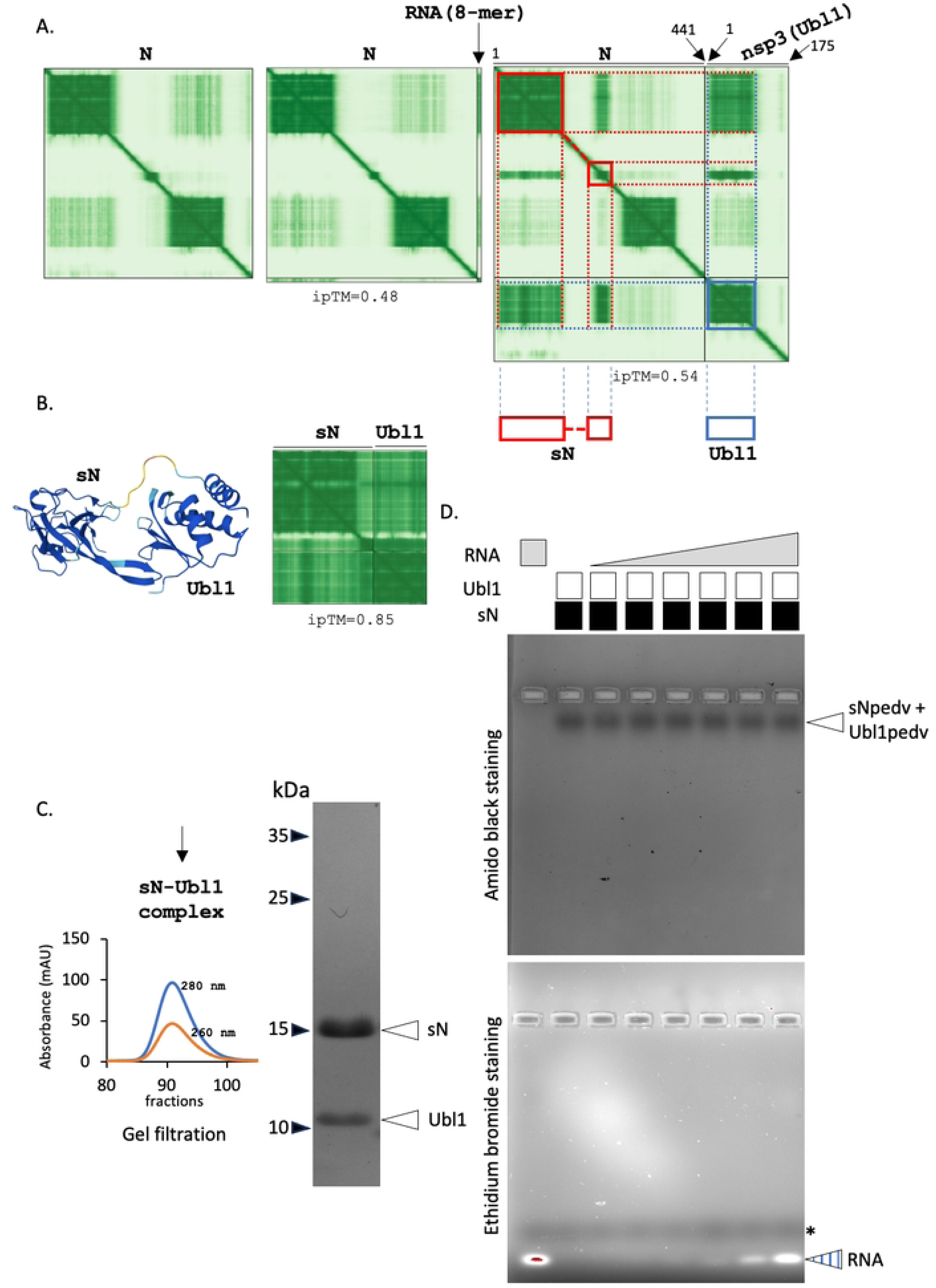
AlphaFold3 predicted structures of the PEDV nucleoprotein N complexed with the N-ter domains of nsp3 (A). AF3 predicted that the Ubl1 domain of nsp3 interacts to N through two distinct domains (N2 and an helix in N3, allowing the design of sN and Ubl1 as the minimal constructs able to bind. The expected position error (in Angströms) of the best predicted structure of the complex is indicated as in Fig. 1. (B) AF3 prediction of the structure of the sN-Ubl1 complex. (C) Final step of the sN-Ubl1 complex purification by gel filtration (left). SDS-PAGE of the purified sN-Ubl1 complex (right). (D) Native agarose gel electrophoresis of the sN - Ubl1 complex and RNA. The gel was stained with Amido black to reveal proteins (upper panel) and with ethidium bromide for nucleic acids detection (lower panel). Bands corresponding to the protein complex and free RNA. RNA was not able to displace nsp3 complexed with N. * indicates the bromophenol blue tracking dye migration.

## Discussion

One of the main features in the coronavirus cycle in the infected cell is the compartmental separation of viral RNA synthesis and nucleocapsid assembly. While the machinery of coronavirus RNA synthesis (that comprises most of the 16 nsps that derive from the proteolytic processing of the two polyproteins pp1a and pp1b) reside inside the virally-induced double-membrane vesicle (DMV) organelles, viral mRNAs including genomic RNA are exported through molecular pores spanning both membrane bilayers. These pores are constituted of nsp4 and nsp3 assemblies, with a large part of nsp3 residing on the cytoplasmic face of the DMV and projecting into the cytoplasm its amino-terminal domain (Wolff et al., 2020). The nucleoprotein N was established to interact only with the nsp3 of the coronavirus RNA synthesis machinery, and in particular, to its Ubl1-Ac N-ter domains. This interaction between N and nsp3 is believed to be critical to ensure that naked genomic RNA that exit from the DMV following synthesis is encapsidated by N bound to nsp3 at the vicinity of the channel exit. This knowledge is essentially based from recent studies, most of which being carried out on the prototypal coronavirus MHV, or in response to the emergence of the SARS-CoV-2 coronavirus responsible for the COVID-19 pandemic, both of them belonging to the *β-coronavirus* genus.

In the present study, we analyzed the interaction play between N, nsp3 and RNA of an a-coronavirus (PEDV) to encapsidate RNA. We found that N was able to form a stable complex with the N-ter extremity of nsp3 with an affinity in the 100 nM - 200 nM range. A short RNA (an 8-nt long RNA) as well as a 60-nt long hairpin RNA were able to fully displace nsp3 from its complex to N and to form a complex with N. Affinity of a small RNA to the N was found to be in the pM range. We also observed that nsp3 was not able to displace the 8-mer RNA when it was complexed with N, suggesting that the interaction play between N, nsp3 and RNA is directional.

Taking our results as a whole, we propose a model (**Fig. 7**) that restrict the interaction play between N, nsp3 and RNA to nucleocapsid assembly when (genomic) RNA exit from the nsp4/nsp3 channel. Thus, small stretches of the RNA that exit from the pore are able to bind to N that was previously recruited by nsp3. RNA binding to the nsp3-complexed N results in the release of nsp3 from N-nsp3 complexes and the assembly of genomic RNA into progeny nucleocapsids in the cytoplasm. The difference of affinity of the N for nsp3 (with a K_D_ around 150nM) and for a 8-mer RNA (K_D_ in the picomolar range) gives strong support to this model. As a consequence, we believe that this K_D_ differential between these three species does not support the idea of a possible dissociation of the ribonucleoprotein complex mediated by nsp3 in the early steps of infection to allow genome entry into DMV as previously proposed for SARS-CoV-2 (Ni et al., 2023).

**Figure 7.**
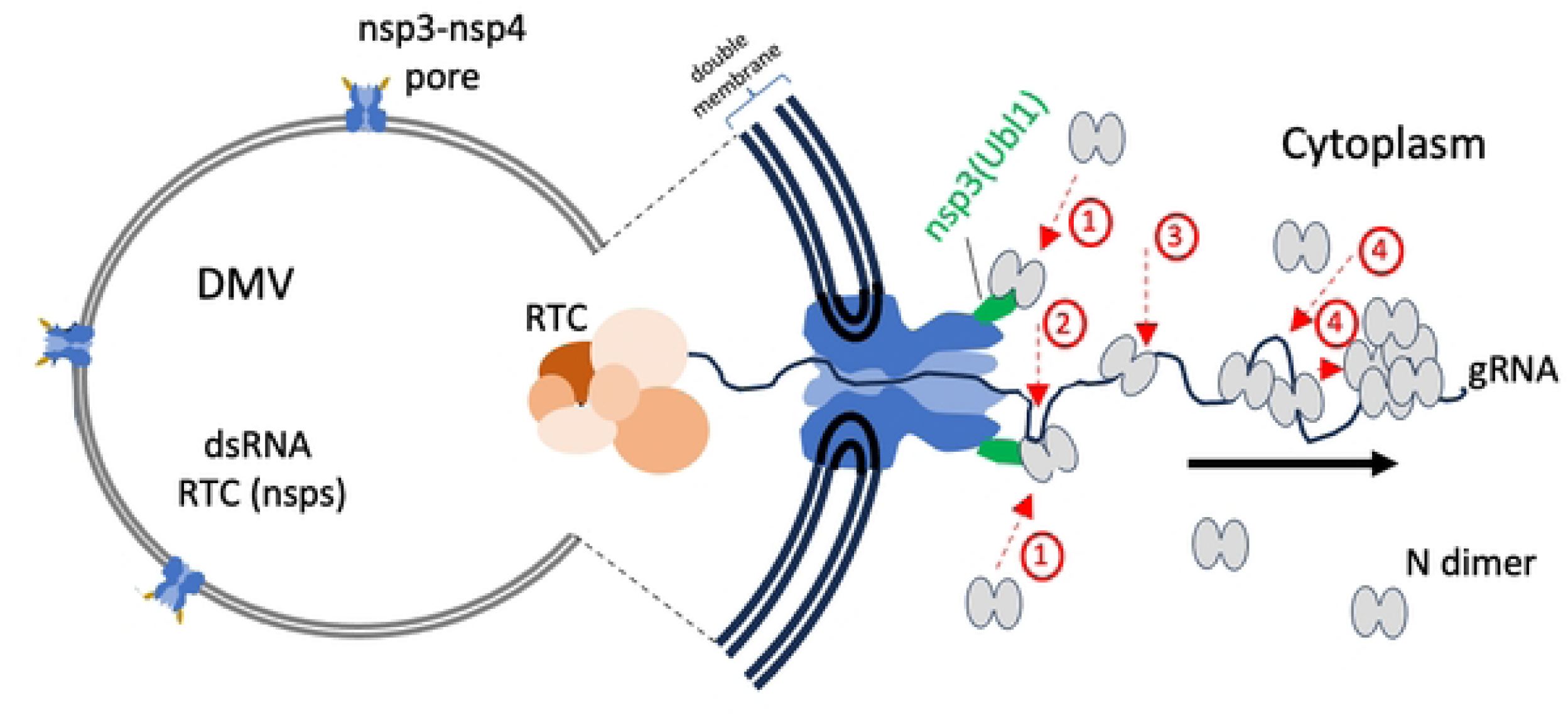
Model of the DMV that contains the coronavirus RTC. Spanning both membranes are multiple copies of the pore, composed of an assembly of nsp3 and nsp4 subunits, which serve as a channel between the DMV interior and the cytoplasm. The amino-terminal Ubl1 domain of nsp3 (nsp3(Ubl1) is shown in green. After its neosynthesis by the RTC in the DMV, gRNA is exported out of the DMV to assemble with N protein previously bound to nsp3(Ubl1) to form progeny ribonucleoprotein complexes in the cytoplasm that will be integrated in virions. (1) N is recruited by nsp3(Ubl1), (2) RNA binds the N-nsp3 complex, (3) RNA is complexed with N and nsp3 is released from the complex, (4) N-RNA complexes oligomerize through N-N interactions.

We also identified a minimal stable complex between N and nsp3 with its Ubl1 domain bound to a N2-3 form of N. Using AF3 structure prediction tool, we designed a truncated form of N that included its N2 (structured) domain fused to its N3 hydrophobic helix (but deleted from its SR and Asn-rich N3 stretches). This sN-Ubl1 complex was found refractory to RNA recruitment, suggesting that RNA binding site(s) on the N nucleoprotein does not strictly overlap the nsp3 binding site(s).

It is interesting to speculate on the importance of the direct and crucial interaction of the N protein with nsp3 that was identified among viruses of the α-and β-coronavirus genera (Masters, 2026 and references therein) on their genetic evolution. The fact that these two proteins that are encoded by two highly distant locus on their genome suggests that during evolution, coronaviruses should preserve their ability to interact. This apparently strongly contrasts with the capacity of coronavirus genome to recombine efficiently, even between genetically distant viruses. As an example, it was thus reported in 2016 the emergence in the European swine population of the emergence of TGEV/PEDV recombinants, sometimes referred as “SeCovs” (Belsham et al. 2016; Boniotti et al., 2016). Examination of the TGEV and PEDV nsp3 and N sequences suggest that these two proteins are not prone to cross-interact. It is worth mentioning that these natural recombinants all display a N-nsp3 couple that derives from the same (TGEV) genome backbone.

All the experiments described in this study were carried out with a (bacterially-expressed) unphosphorylated N protein form of PEDV that appears fully competent for binding to nsp3 and (small) RNA species. It should be noted here that for numerous β-coronaviruses and PEDV, the intracellular N protein is hyperphosphorylated and the N present in virions is hypo-(or not) phosphorylated (Hogue, 1995; Wu et al., 2009, Jia et al., 2025). It was recently shown on SARS-CoV-2 that N phosphorylation events coordinated by three host protein kinases results in complete inhibition of RNA binding (Botova et al., 2024), thus suggesting that recruitment of specific host kinases (and possibly phosphatase(s)?) control the ability of N to bind nsp3 and RNA. It should be of interest in the future to identify on PEDV the phosphorylated residues of N in infected cells, the kinases associated to these phosphorylation events and to determine the extent to which phosphorylation controls and regulates the recruitment of nsp3 and RNA by the N protein to regulate genome encapsidation.

In conclusion, we found that the N-nsp3 interaction that was identified in several β-coronaviruses was preserved in an α-coronavirus. We determined the affinities of N for nsp3 and of the N to a short RNA stretch, affinities that display a difference of two orders of magnitude in favor of the N-RNA interaction. We observed that while a small RNA is able to displace nsp3 complexed to N to form a RNA-N complex, while nsp3 was not able to bind and to displace RNA complexed to the N protein. Using a minimal N-nsp3 complex, we also evidenced that nsp3 and RNA binding sites are distinct on N. We proposed a model for viral RNA assembly through a primary interaction between N and nsp3 at the cytoplasmic face of the DMV.

## Materials and Methods

### Bacterial Expression and Purification of recombinant Proteins

Recombinant proteins were produced in *E. coli* BL21 bacteria. Protein sequences are shown in **Fig. S1**. The N recombinant protein contains a 6xHis-tag at its N-terminus and a TEV cleavage site and was expressed using the pET28a+ plasmid. The nsp3(Ubl1) domain was expressed as a GST-fusion protein with a TEV cleavage site using the pGEX4.1 vector as previously described (Galloux et al., 2012). BL21 bacteria precultures (4ml) were grown overnight at 37^◦^C in Luria-Bertani (LB) medium supplemented with 50 μg/mL kanamycin for N expression or 100 μg/mL ampicillin for GST-nsp3(Ubl1) expression. Then, 500 ml of antibiotic-supplemented LB medium were seeded with 200 μl of precultures during 8 hrs. Next, an equivalent volume of LB was added to the culture, and protein expression was induced with 330 μM isopropyl-β-D-thio-galactoside (IPTG) for 15 h at 28^◦^ C. Bacteria were then harvested by centrifugation (5,000 x rpm, 15 min, rotor TA-10-250 Beckman Coulter).

#### GST-nsp3(Ubl1) purification

The bacterial pellet was resuspended in 40 ml of lysis buffer (50 mM Tris-HCl pH 7.5, 60 mM NaCl, 1 mM EDTA, 2 mM dithiothreitol (DTT), 0.2% Triton X-100, 1 mg/mL lysozyme). A pellet of the complete protease inhibitor cocktail (Roche) was added to the suspension for an incubation of 1 h on ice. Next, suspensions were sonicated using a Qsonica sonicator (two times 1 min with an amplitude signal of 40), and 4 μL of benzonase (250 U/µl, UGAP) was added for a 30 min period at room temperature. Next, NaCl was added to reach a 1M final concentration during 30 min, next, the lysate was centrifugated at 4^◦^C for 30 min at 10,000 × rpm (rotor TA-10-250, Beckman-Coulter). Glutathione-Sepharose 4B beads (1.5 ml, Cytiva) were added to the clarified supernatant and incubated at 4^◦^C for a 3hrs to overnight period. Next, the beads were washed once in 1M NaCl lysis buffer, two times in 20 mM Tris-HCl pH 7.5, 1M NaCl and once in 20 mM Tris-HCl pH 7.5, 150 mM NaCl. Next, beads were incubated with the TEV protease (6µl, Sigma-Aldrich) in 20mM Tris-HCl pH7.5, 150mM NaCl, 0.5mM EDTA, 1mM DTT buffer for 48 hrs at room temperature. The supernatant was then loaded onto a Superdex 200 16/30 column (GE Healthcare) buffered with 20 mM Tris-HCl pH 7.5, 500 mM NaCl. Fractions containing the nsp3(Ubl1) domain were pooled and protein concentration was determined with a Bradford assay (Biorad) calibrated with BSA.

#### His-tagged N purification

The bacterial pellet was resuspended in 40 ml of lysis buffer (20 mM Tris-HCl pH 7.5, 500 mM NaCl, 0.1% Triton X-100, 10mM imidazole, 1 mg/mL lysozyme). A pellet of the complete protease inhibitor cocktail (Roche) was added to the suspension for an incubation of 1 h on ice. Next, suspensions were sonicated using a Qsonica sonicator (two times 1 min with an amplitude signal of 40), and 4µl of benzonase (250U/µl, UGAP) was added. Next, NaCl was added to reach a 1M final concentration during 30 min, next, the lysate was centrifugated at 4^◦^C for 30 min at 10,000 × rpm (rotor TA-10-250, Beckman Coulter). Chelatine sepharose fast flow beads (Cytiva) saturated with NiSO4 were added to the clarified supernatant and incubated at room temperature for 1 hour. Next, beads were washed twice in 1M NaCl lysis buffer and successive incubations in 20mM Tris-HCl pH7.5, 1M NaCl with increasing concentrations of imidazole (10mM, 50mM to 100mM). Next, N protein was eluted with 20mM Tris-HCl, 1M NaCl, 800mM imidazole. Proteins were then loaded onto a Superdex 200 16/30 column (GE Healthcare) buffered with 20mM Tris-HCl pH7.5, 500mM NaCl buffer. N containing fractions were pooled and protein concentration was determined with a Bradford assay (Biorad).

#### nsp3(Ubl1)-N complex production and purification

Cultures of BL21 bacteria harboring the pET28a+ and the pGEX4.1 derivates were grown overnight at 37^◦^C in Luria-Bertani (LB) medium supplemented with 50 μg/mL kanamycin and 100 μg/mL ampicillin. Then, 500 ml of antibiotic-supplemented LB medium were seeded with 200 μl of precultures during 8 hrs. Next, an equivalent volume of LB was added to the culture, and protein expression was induced with 330 μM isopropyl-β-D-thio-galactoside (IPTG) for 15 h at 28 ◦C. Bacteria were then harvested by centrifugation (5,000 x rpm, 15 min, rotor TA-10-250 Beckman Coulter). The bacterial pellet was resuspended in 40 ml of lysis buffer (50 mM Tris-HCl pH 7.5, 60 mM NaCl, 0.2% Triton X-100, 1 mg/mL lysozyme) and processed for GST-chromatography followed by a TEV protease treatment and gel filtration as described above. Fractions containing the nsp3(Ubl1)-N complexes were pooled and protein concentration was determined with a Bradford assay (Biorad).

### Affinity determination by Bio layer interferometry

Binding kinetics experiments were performed on an Octet system (Octet RED96) (FortéBio, CA). A black bottom 96-well microplate (Greiner Bio-One # 655209) was filled with 200 uL of solution (N or nsp3(Ubl1) in 20 mM Tris pH 7.5, 500 mM NaCl) and agitated at 1 000 rpm, and all experiments were carried out at 25°C. Tips were hydrated in the Tris buffer for 1 hour at room temperature prior experiments. Biotinylated RNA (concentration?) or His-tagged N (x µg/mL) were loaded on streptavidin SA (18-0009) or NTA biosensors (Pall ForteBio) for 1 min, respectively. After a baseline step in assay buffer (20 mM Tris pH 7.5, 500 mM NaCl, 0.1% bovine serum albumin, 0.02% Tween 20) and a quenching step in 5 ug/mL biocytin, ligand-loaded sensors were dipped into known concentrations of protein for an association phase during 500 seconds. The sensors were then dipped in assay buffer for a dissociation step during 500 seconds in assay buffer. Association and dissociation curves were globally fitted to a 1:1 binding model. Binding curves were fit using the “association then dissociation” equation in the FortéBio Data analysis software version 7.1 to calculate KD.

### Native gel electrophoresis

Proteins and RNAs were mixed in a 20 μL final volume 15 min at 4°C before electrophoresis analysis. The sucrose loading buffer was added to the samples for migration on a native 1% agarose gel containing ethidium bromide (0.5 μg/ml). Migration was performed in 1× Tris– glycine buffer during 1 hour and 15 min at 80V, before analyzing RNA migration using a UV plate reader. The gels were then stained with Amido black 10B before being decolorized with 5% acetic acid.

## Acknowledgements

The present work has benefited from the I2BC’s Macromolecular Interactions Measurement Platform (MIP) Core facility, (Paris-Saclay University) supported by French Infrastructure for Integrated Structural Biology (FRISBI) ANR-10-INBS-05.

## Funding and additional information

B.D. acknowledges support of the ANR-17-CE18-0006-01 program. The funders had no role in study design, data collection and analysis, decision to publish, or preparation of the manuscript.

## References

Anderson TK, Hoferle PJ, Chojnacki KJ, Lee KW, Coon JJ, Kirchdoerfer RN. An alphacoronavirus polymerase structure reveals conserved replication factor functions. Nucleic Acids Res. 2024 Jun 10;52(10):5975–5986.

Belsham GJ, Rasmussen TB, Normann P, Vaclavek P, Strandbygaard B, Bøtner A. Characterization of a Novel Chimeric Swine Enteric Coronavirus from Diseased Pigs in Central Eastern Europe in 2016. Transbound Emerg Dis. 2016 Dec;63(6):595–601.

Bessa LM, Guseva S, Camacho-Zarco AR, Salvi N, Maurin D, Perez LM, Botova M, Malki A, Nanao M, Jensen MR, Ruigrok RWH, Blackledge M. The intrinsically disordered SARS-CoV-2 nucleoprotein in dynamic complex with its viral partner nsp3a. Sci Adv. 2022 Jan 21;8(3):eabm4034.

Boniotti MB, Papetti A, Lavazza A, Alborali G, Sozzi E, Chiapponi C, Faccini S, Bonilauri P, Cordioli P, Marthaler D. Porcine Epidemic Diarrhea Virus and Discovery of a Recombinant Swine Enteric Coronavirus, Italy. Emerg Infect Dis. 2016 Jan;22(1):83–7.

Botova M, Camacho-Zarco AR, Tognetti J, Bessa LM, Guseva S, Mikkola E, Salvi N, Maurin D, Herrmann T, Blackledge M. A specific phosphorylation-dependent conformational switch in SARS-CoV-2 nucleocapsid protein inhibits RNA binding. Sci Adv. 2024 Aug 2;10(31):eaax2323.

Galloux M, Tarus B, Blazevic I, Fix J, Duquerroy S, Eléouët JF. Characterization of a viral phosphoprotein binding site on the surface of the respiratory syncytial nucleoprotein. J Virol. 2012 Aug;86(16):8375–87.

Ghosh S, Dellibovi-Ragheb TA, Kerviel A, Pak E, Qiu Q, Fisher M, Takvorian PM, Bleck C, Hsu VW, Fehr AR, Perlman S, Achar SR, Straus MR, Whittaker GR, de Haan CAM, Kehrl J, Altan-Bonnet G, Altan-Bonnet N. β-Coronaviruses Use Lysosomes for Egress Instead of the Biosynthetic Secretory Pathway. Cell. 2020 Dec 10;183(6):1520–1535.e14.

Cong Y, Ulasli M, Schepers H, Mauthe M, V’kovski P, Kriegenburg F, Thiel V, de Haan CAM, Reggiori F. Nucleocapsid Protein Recruitment to Replication-Transcription Complexes Plays a Crucial Role in Coronaviral Life Cycle. J Virol. 2020 Jan 31;94(4):e01925–19.

Hogue BG. Bovine coronavirus nucleocapsid protein processing and assembly. Adv Exp Med Biol. 1995;380:259–63.

Huang Y, Wang T, Zhong L, Zhang W, Zhang Y, Yu X, Yuan S, Ni T. Molecular architecture of coronavirus double-membrane vesicle pore complex. Nature. 2024 Sep;633(8028):224–231.

Hurst KR, Ye R, Goebel SJ, Jayaraman P, Masters PS. An interaction between the nucleocapsid protein and a component of the replicase-transcriptase complex is crucial for the infectivity of coronavirus genomic RNA. J Virol. 2010 Oct;84(19):10276–88.

Hurst KR, Koetzner CA, Masters PS. Characterization of a critical interaction between the coronavirus nucleocapsid protein and nonstructural protein 3 of the viral replicase-transcriptase complex. J Virol. 2013 Aug;87(16):9159–72.

Jia X, Chen J, Li C, Li J, Su M, Yang K, Zhang Y, Li Z. Glycogen synthase kinase 3 promotes the proliferation of porcine epidemic diarrhoea virus by phosphorylating the nucleocapsid protein Virulence. 2025 Dec;16(1):2506504.

Klumperman J, Locker JK, Meijer A, Horzinek MC, Geuze HJ, Rottier PJ. Coronavirus M proteins accumulate in the Golgi complex beyond the site of virion budding. J Virol. 1994 Oct;68(10):6523–34.

Krijnse-Locker J, Ericsson M, Rottier PJ, Griffiths G. Characterization of the budding compartment of mouse hepatitis virus: evidence that transport from the RER to the Golgi complex requires only one vesicular transport step. J Cell Biol. 1994 Jan;124(1-2):55–70.

Lei J, Kusov Y, Hilgenfeld R. Nsp3 of coronaviruses: Structures and functions of a large multi-domain protein. Antiviral Res. 2018 Jan;149:58–74.

Masters PS. Coronavirus genome packaging and nucleocapsid assembly. J Virol. 2026 Feb 17;100(2):e0133025.

McBride R, van Zyl M, Fielding BC. The coronavirus nucleocapsid is a multifunctional protein Viruses. 2014 Aug 7;6(8):2991–3018.

Millet JK, Jaimes JA, Whittaker GR. Molecular diversity of coronavirus host cell entry receptors. FEMS Microbiol Rev. 2021 May 5;45(3):fuaa057.

Neuman BW, Angelini MM, Buchmeier MJ. Does form meet function in the coronavirus replicative organelle? Trends Microbiol. 2014 Nov;22(11):642–7.

Neuman BW. Bioinformatics and functional analyses of coronavirus nonstructural proteins involved in the formation of replicative organelles. Antiviral Res. 2016 Nov;135:97–107.

Ni X, Han Y, Zhou R, Zhou Y, Lei J. Structural insights into ribonucleoprotein dissociation by nucleocapsid protein interacting with non-structural protein 3 in SARS-CoV-2. Commun Biol. 2023 Feb 18;6(1):193.

Snijder EJ, Limpens RWAL, de Wilde AH, de Jong AWM, Zevenhoven-Dobbe JC, Maier HJ, Faas FFGA, Koster AJ, Bárcena M. A unifying structural and functional model of the coronavirus replication organelle: Tracking down RNA synthesis. PLoS Biol. 2020 Jun 8;18(6):e3000715.

V’kovski P, Kratzel A, Steiner S, Stalder H, Thiel V. Coronavirus biology and replication: implications for SARS-CoV-2. Nat Rev Microbiol. 2021 Mar;19(3):155–170.

Wang Q, Vlasova AN, Kenney SP, Saif LJ. Emerging and re-emerging coronaviruses in pigs. Curr Opin Virol. 2019 Feb;34:39–49.

Wang Q, Wu J, Wang H, Gao Y, Liu Q, Mu A, Ji W, Yan L, Zhu Y, Zhu C, Fang X, Yang X, Huang Y, Gao H, Liu F, Ge J, Sun Q, Yang X, Xu W, Liu Z, Yang H, Lou Z, Jiang B, Guddat LW, Gong P, Rao Z. Structural Basis for RNA Replication by the SARS-CoV-2 Polymerase. Cell. 2020 Jul 23;182(2):417–428.e13.

Wolff G, Limpens RWAL, Zevenhoven-Dobbe JC, Laugks U, Zheng S, de Jong AWM, Koning RI, Agard DA, Grünewald K, Koster AJ, Snijder EJ, Bárcena M. A molecular pore spans the double membrane of the coronavirus replication organelle. Science. 2020 Sep 11;369(6509):1395–1398.

Wu CH, Yeh SH, Tsay YG, Shieh YH, Kao CL, Chen YS, Wang SH, Kuo TJ, Chen DS, Chen PJ. Glycogen synthase kinase-3 regulates the phosphorylation of severe acute respiratory syndrome coronavirus nucleocapsid protein and viral replication J Biol Chem. 2009 Feb 20;284(8):5229-39.

Yan L, Ge J, Zheng L, Zhang Y, Gao Y, Wang T, Huang Y, Yang Y, Gao S, Li M, Liu Z, Wang H, Li Y, Chen Y, Guddat LW, Wang Q, Rao Z, Lou Z. Cryo-EM Structure of an Extended SARS-CoV-2 Replication and Transcription Complex Reveals an Intermediate State in Cap Synthesis. Cell. 2021 Jan 7;184(1):184–193.e10.

Yan L, Yang Y, Li M, Zhang Y, Zheng L, Ge J, Huang YC, Liu Z, Wang T, Gao S, Zhang R, Huang YY, Guddat LW, Gao Y, Rao Z, Lou Z. Coupling of N7-methyltransferase and 3’-5’ exoribonuclease with SARS-CoV-2 polymerase reveals mechanisms for capping and proofreading. Cell. 2021 Jun 24;184(13):3474–3485.e11.

Zargarian L, Ai X, Song ZY, Perez-Gorgol L, Couprie-Diaz E, Fossé P, Xi XG, Mauffret O. Chaperone Activity of SARS-CoV-2 Nucleocapsid Protein: RNA Annealing and Destabilization Mechanisms. J Mol Biol. 2025 Oct 1;437(19):169312.

Zúñiga S, Sola I, Moreno JL, Sabella P, Plana-Durán J, Enjuanes L. Coronavirus nucleocapsid protein is an RNA chaperone. Virology. 2007 Jan 20;357(2):215–27.

